# Gaussian curvature dilutes the nuclear lamina, favoring nuclear rupture, especially at high strain rate

**DOI:** 10.1101/2021.10.13.464257

**Authors:** Charlotte R. Pfeifer, Michael P. Tobin, Sangkyun Cho, Manasvita Vashisth, Larry Dooling, Lizeth Lopez Vazquez, Emma G. Ricci-De Lucca, Keiann T. Simon, Dennis E. Discher

## Abstract

Nuclear rupture has long been associated with deficits or defects in lamins, with recent results also indicating a role for actomyosin stress, but key physical determinants of rupture remain unclear. Here, lamin-B stably interacts with the nuclear membrane at sites of low Gaussian curvature yet dilutes at high-curvature to favor rupture, whereas lamin-A depletes similarly but only at high strain-rates. Live cell imaging of lamin-B1 gene-edited cancer cells is complemented by fixed-cell imaging of ruptured nuclei in: iPS-derived cells from progeria patients, cells within beating chick embryo hearts, and cancer cells that develop multiple ruptures in migrating through small pores. Dilution and curvature-dependent rupture fit a parsimonious model of a stiff filament that detaches from a curved surface, suggesting an elastic-type response of lamin-B, but rupture is also modestly suppressed by inhibiting myosin-II and by hypotonic stress, which slow the strain rates. Lamin-A dilution and nuclear rupture likelihood indeed increase above a threshold rate of pulling into small pipettes, suggesting a viscoplastic coupling to the envelope for protection against nuclear rupture.

**Summary statement:** High nuclear curvature drives lamina dilution and nuclear envelope rupture even when myosin stress is inhibited. Stiff filaments generally dilute from sites of high Gaussian curvature, providing mathematical fits of experiments.

## Introduction

During interphase, the nuclear envelope typically functions as a barrier between the nucleus and cytoplasm. Rupture of the envelope mis-localizes key nuclear proteins to cytoplasm, including transcription factors (De Vos et al., 2011) and DNA repair factors (Irianto et al., 2016; Irianto et al., 2017), as well as nuclear entry of DNA-binding proteins (Denais et al., 2016; Irianto et al., 2017; Maciejowski et al., 2015; Raab et al., 2016). Mechanisms of rupture continue to be elucidated, especially biophysical determinants, but early work associated rupture with deficiencies or defects in lamins (Hatch et al., 2013; Tamiello et al., 2013; Vargas et al., 2012; Vergnes et al., 2004). A- and B-type lamins make ~½-μm long filaments that resist bending within juxtaposed meshworks at the inner nuclear membrane (Turgay et al., 2017), and farnesylated lamin-B1 and -B2 associate more directly with the membrane than non-farneslyated lamin-A,C (**Fig.1A**). Lamin-B levels are nearly constant across different tissues (Swift et al., 2013) and double in level as a cell duplicates its DNA (Vashisth et al., 2021), whereas lamin-A,C increases from low levels in soft embryos and brain to high levels in stiff muscle and rigid bone (Cho et al., 2019; Swift et al., 2013). Various cells studied here are representative with lamin-A:B ratios that range from ~1:1 in early chick embryo hearts (Cho et al., 2019) to ~2:1 in A549 human lung cancer cells (Harada et al., 2014) and ~7:1 in mesenchymal stem/progenitors derived from progeria patient iPS cells (Cho et al., 2018) – all of which provide broad insight into how the lamins confer nuclear strength and stability.

**Figure 1.**
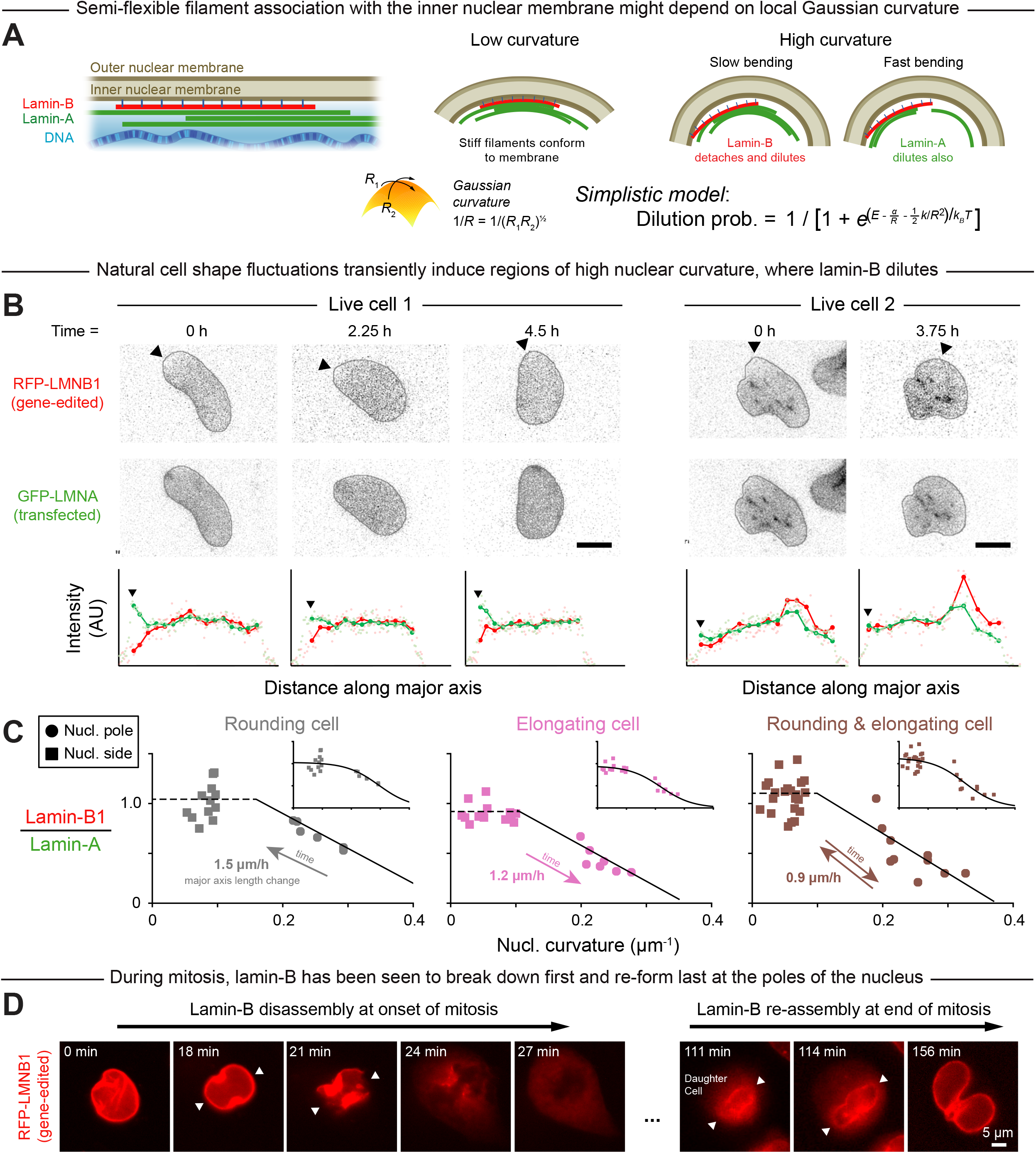
High Gaussian curvature disrupts the nuclear lamina. A. A-type and B-type lamins form juxtaposed networks inside the nuclear membrane. Lamin filaments are expected to stably interact with a membrane of low Gaussian curvature but dissociate from a high-curvature membrane (“Single-filament model,” Materials & Methods). *R* = Gaussian radius of curvature; *R*_1_, *R*_2_ = principal curvature radii. Lamin-A and lamin-B networks are thought to have differing material properties—and hence to dilute, or be depleted, at differing rates from regions of high nuclear curvature. B. Images: A549 human lung cancer cells with RFP-lamin-B1 from monoallelic gene editing were transfected with GFP-lamin-A and live-imaged over several hours. Cell shape fluctuations transiently induced regions of high nuclear curvature. Intensity profiles: measured along the cell’s major axis, originating at the arrow head. Representative of 10 cells/20 nuclear poles; scale = 10 *μ*m. C. Lamin-A and -B intensities were measured at the poles and sides of nuclei in three representative cells that rounded and/or elongated during live-imaging. Plots show lamin-B to lamin-A intensity ratio as a function of nuclear curvature. Insets: same data, fit with a sigmoidal function per our simple filament (un)binding model. Representative of 4 cells/8 nuclear poles. D. Live images of A549 mitosis, with the lamin-B network disassembling first and reassembling last at the higher-curvature poles of the nucleus (arrow heads).

Tension in the nuclear membrane might cause rupture and could result from pressure differences and other forces (Zhang et al., 2019). Continuity of the outer nuclear membrane with the endoplasmic reticulum would tend to dissipate high tensions and oppose nuclear membrane rupture, which suggests additional mechanisms. High-curvature AFM tips cause nuclear-specific rupture when pressed on nuclei within intact adherent cells (Xia et al., 2018), and high-curvature micronuclei are also most susceptible to rupture (Xia et al., 2019). Such observations have thus associated nuclear envelope rupture with positive Gaussian curvature as opposed to simple mean curvature (**Fig.S1A**). Relationships otherwise remain unclear between site or sites of nuclear rupture and nuclear curvature, including negative Gaussian curvature (saddle shapes), and time- or rate-dependent density changes of A- and B-lamins. Distinctive effects might be expected for the lamin-A and lamin-B networks because they respectively exhibit viscoplastic or elastic behavior (Swift et al., 2013). We hypothesized that the elastic lamin-B meshwork would dilute comparatively quickly in regions of high curvature, whereas the more viscoplastic lamin-A would depend more strongly on the rate of change of nuclear curvature (Fig.1A).

## Results and discussion

### Dilution of lamin-B at nuclear poles fits a simple equilibrium model

We first imaged local changes in lamin levels as nuclear curvature fluctuated over hours within sparsely plated A549 cells that spread, elongated, and/or rounded-up in standard culture. The cells were gene-edited to express RFP-lamin-B1 in order to minimize strong effects of lamin-B1 perturbations on cell cycle (Shimi et al., 2011; Vergnes et al., 2004), and they were also transfected with GFP-lamin-A for live-imaging. Lamin-B1 was often diluted in regions of high curvature unlike lamin-A (**Fig.1B**). Regardless of the quasi-static nuclear extension or rounding, the lamin B-to-A intensity ratio was low at the nuclear poles but not the flat sides or other low-curvature regions (**Fig.1C**). Immunostaining after fixation of distinct A549 clones with persistent ‘spindle-shaped’ nuclei confirmed the relative depletion of lamin-B at high-curvature nuclear poles, as did imaging of U2OS human bone cancer cells (**Fig.S1B-D**).

Live-imaging results for curvature-dependent lamin-B depletion fit a parsimonious, ‘spherical cow’-type model (“Single-filament model,” Materials & Methods) based on two ideas. First, semi-flexible lamin-B filaments stably bind to an inner nuclear membrane of low Gaussian curvature but quickly detach from a high-curvature membrane. Second, the lamin-A layer dilutes minimally at the low deformation rates typical of cell and nuclear movements in 2D culture.

Mitotic cells further show the lamin-B meshwork disassembled first at the highest curvature nuclear poles and also re-assembled last at such sites (**Fig.1D**, **Fig.S2**). Nuclear curvature might thus contribute to lamin-B disassembly-reassembly in mitosis together with other biophysical and established molecular mechanisms (Beaudouin et al., 2002; Salina et al., 2002). Whereas holes in the lamin-B network are typical of the mitotic initiation of nuclear envelope breakdown (Collas, 1998; Georgatos et al., 1997), local depletion of lamin-B in interphase nuclei can lead to aberrant nuclear rupture.

### Rupture at nuclear poles in progeria-derived cells and within chick embryos

As a second model, mesenchymal stem/progenitor cells were differentiated from iPS cells with mutant lamin-A from patients with Hutchinson-Gilford progeria syndrome (De Sandre-Giovannoli, 2003; Eriksson et al., 2003) because such defects should favor nuclear rupture in cultures on rigid substrates that promote actomyosin stress in spreading of the cell and its nucleus (Tamiello et al., 2013; Xia et al., 2018). Frequent rupture was indeed indicated by formation of lamin-B-deficient blebs in immunofluorescence (**Fig.2A**). Importantly, the lamin-A containing blebs again occurred overwhelmingly at the poles of the nucleus (**Fig.2B**), confirming that high curvature correlates not only with lamin-B dilution, but further with nuclear envelope rupture.

**Figure 2.**
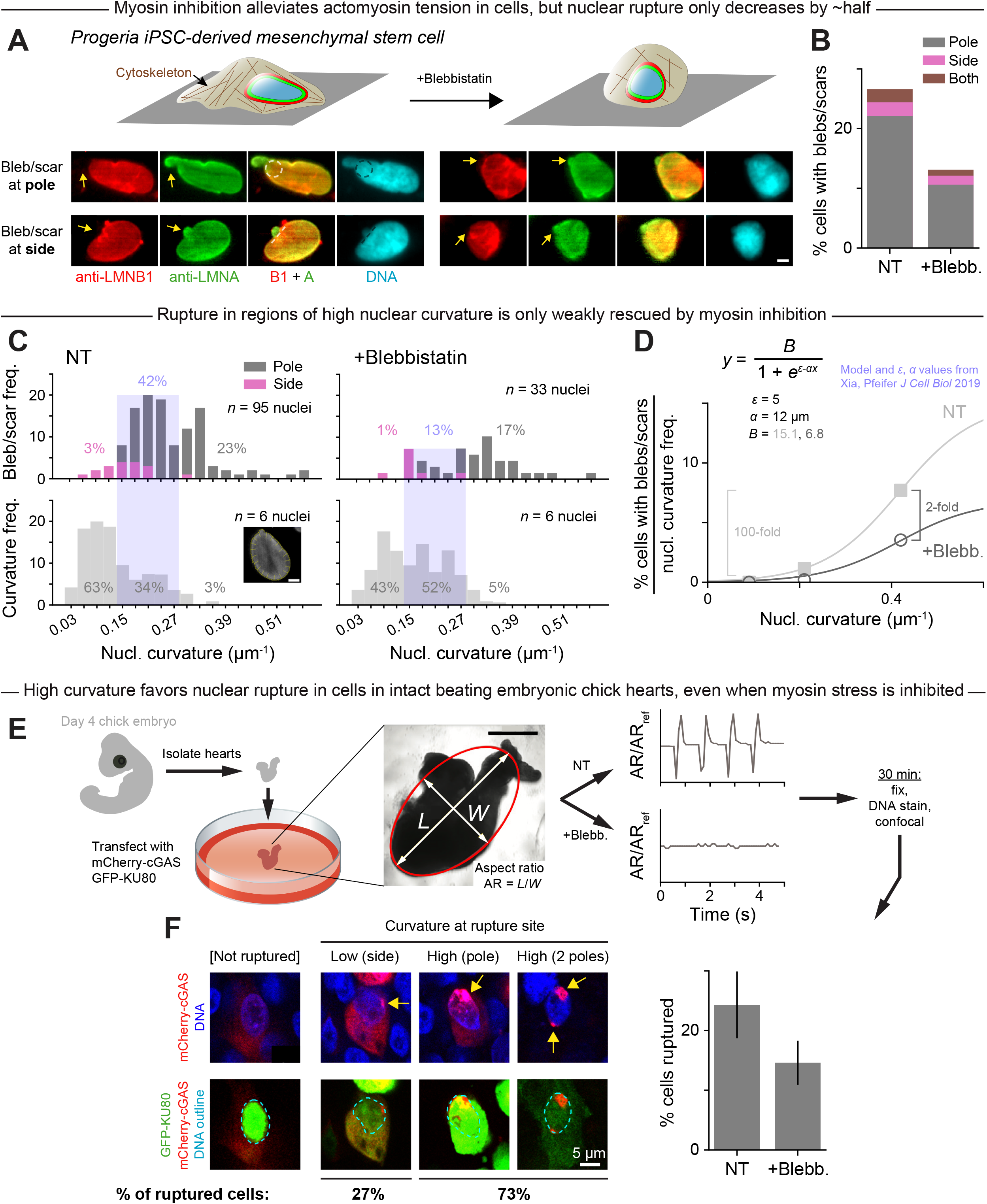
In progeria cells and cells in intact beating embryonic chick hearts, high Gaussian curvature favors nuclear rupture, even when myosin stress is inhibited. A. Mesenchymal stem/progenitor cells from patients with progeria have a defective nuclear lamina, which increases nuclear rupture in standard 2D culture. The nuclear bleb or scar is shows abundant lamin-A but loss of lamin-B (yellow arrows). Myosin-II inhibitor blebbistatin reduces stress on the nucleus, but blebs/scars persist. iPSC = induced pluripotent stem cell. Images: dashed circles and lines trace the local curvature; scale = 5 *μ*m. B. Rupture frequency, as indicated by %progeria cells with blebs/scars, is highest at the nuclear pole and decreases by ~half after blebbistatin (blebb.) treatment. NT = non-treated; 594 cells; ≥242 cells per condition. C. Lower: For 6 non-treated and 6 blebb-treated progeria cells (randomly selected), curvature was measured at ~20 locations around the nuclear perimeter (inset image, scale = 5 *μ*m). Plots show the curvature distributions. Upper: For progeria cells from panel B with blebs/scars, nuclear curvature at the location of the bleb/scar was measured, with blue shading for low, high, or very high curvature. The latter two are enriched for blebs/scars. D. For each of the three nuclear curvature regimes from panel C (low, high, very high), the frequency at which a rupture-inducing curvature falls in that regime (panel C upper) was normalized to the frequency at which that regime occurs in progeria cell nuclei (panel C lower)—and then plotted against the median curvature for that regime. Sigmoidal fit parameters come from our simple filament (un)binding model. E. Hearts isolated from chick embryos were transfected with the nuclear protein GFP-KU80 and the cytoplasmic DNA-binding protein mCherry-cGAS, and then treated with blebb. or control DMSO. During the 30-min live-imaging that followed, blebb. suppressed beating of the heart aspect ratio AR_ref_. *L*, *W* = length, width. Representative of 6 hearts; scale = 200 *μ*m. F. Isolated hearts from panel E were fixed, DNA stained, and imaged. Some heart cells showed nuclear envelope rupture (bar plot) based on nuclear accumulation of cGAS (upper images: yellow arrows) and mis-localization of KU80 outside the nucleus (lower images: dashed outlines indicate nucleus). 72 cells; ≥18 cells per condition; mean ± SEM.

To determine whether rupture is driven by curvature itself or by the actomyosin-generated tensions, we inhibited myosin II and observed rounding of the cells and their nuclei (Fig.2A). Yet rupture frequency decreased by only ~half (Fig.2B) and only by ~one-quarter in the regions of highest nuclear curvature (**Fig.2C**). The results imply curvature-associated rupture occurs largely independent of myosin stress. In contrast to the modest decrease with myosin inhibition, rupture was virtually eliminated in the large regions of low nuclear curvature in both wild-type and blebbistatin-treated progeria cells (**Fig.2D**). This strong dependence of rupture on curvature again fits our simple model of a semi-flexible lamin filament that binds or not to a highly curved nuclear surface (Fig.2D). The results reinforce the idea that curvature-induced lamin-B loss precipitates nuclear envelope rupture.

To address whether curvature-associated rupture also occurs in intact tissues, beating hearts were isolated from day-4 chick embryos, transfected with two factors that reveal nuclear rupture (Cho et al., 2019), treated or not with blebbistatin, and then fixed, DNA-stained, and imaged by confocal microscopy (**Fig.2E**). Myosin inhibition stopped the beating but again caused only a modest decrease in nuclear rupture, with the rupture occurring mostly at the nuclear poles (**Fig.2F**). Rupture was assessed by the cytoplasmic presence of the DNA repair factor GFP-KU80 in the same cells that also show focal nuclear entry of DNA-binding protein mCherry-cGAS. Intact tissue results thus point once again to high curvature, more so than myosin stress, as a predominant driver of nuclear rupture.

### Multiple ruptures in the same nucleus again fit a poleward bias

Some nuclei in the hearts showed multiple sites of rupture (Fig.2F) as was also the case for the progeria cells, suggesting that rupture can occur in a nucleus with a pre-existing bleb. Phospholipid giant vesicles studied in viscous media by Brochard-Wyart and colleagues (Karatekin et al., 2003; Sandre et al., 1999) likewise showed multiple sites of rupture, and each rupture resolved before the next, indicating that high internal pressure caused a vesicle to develop a hole and leak fluid, alleviating pressure, allowing the hole to reseal, but then the cycle repeated with dozens of transient holes in succession. Even without rupture, plasma membrane bleb formation in fibroblasts has been shown to significantly relax intracellular pressure, which only returns to its initial high value once the bleb has retracted (Tinevez et al., 2009). If nuclear rupture is thus primarily driven by intranuclear pressure that is reduced upon rupture, than rupture events are *not* independent. If, however, rupture is primarily driven by curvature, then the persistence of one bleb should be independent of further rupture-blebs at additional high-curvature sites.

To assess independence of multi-site nuclear rupture, painstaking statistics of the number of lamin-B deficient blebs or scars on U2OS nuclei were made after cell migration through constricting pores of different diameters, either custom-made or commercially available (**Fig.3A,B**). Such migration has been seen to squeeze out nucleoplasm (Irianto MBoC 2016), which implies a build-up of nuclear pressure. A single rupture again follows the simple sigmoidal probability *P*_rupt_ as a function of pore-imposed curvature (Fig.1A, Eqn), and — more importantly — the fraction of nuclei with either two ruptures or three or more ruptures fit respectively to (*P*_rupt_)^2^ or (*P*_rupt_)^3^ (**Fig.3C**). These findings imply independent events and argue that rupture is driven by curvature rather than internal pressure.

**Figure 3.**
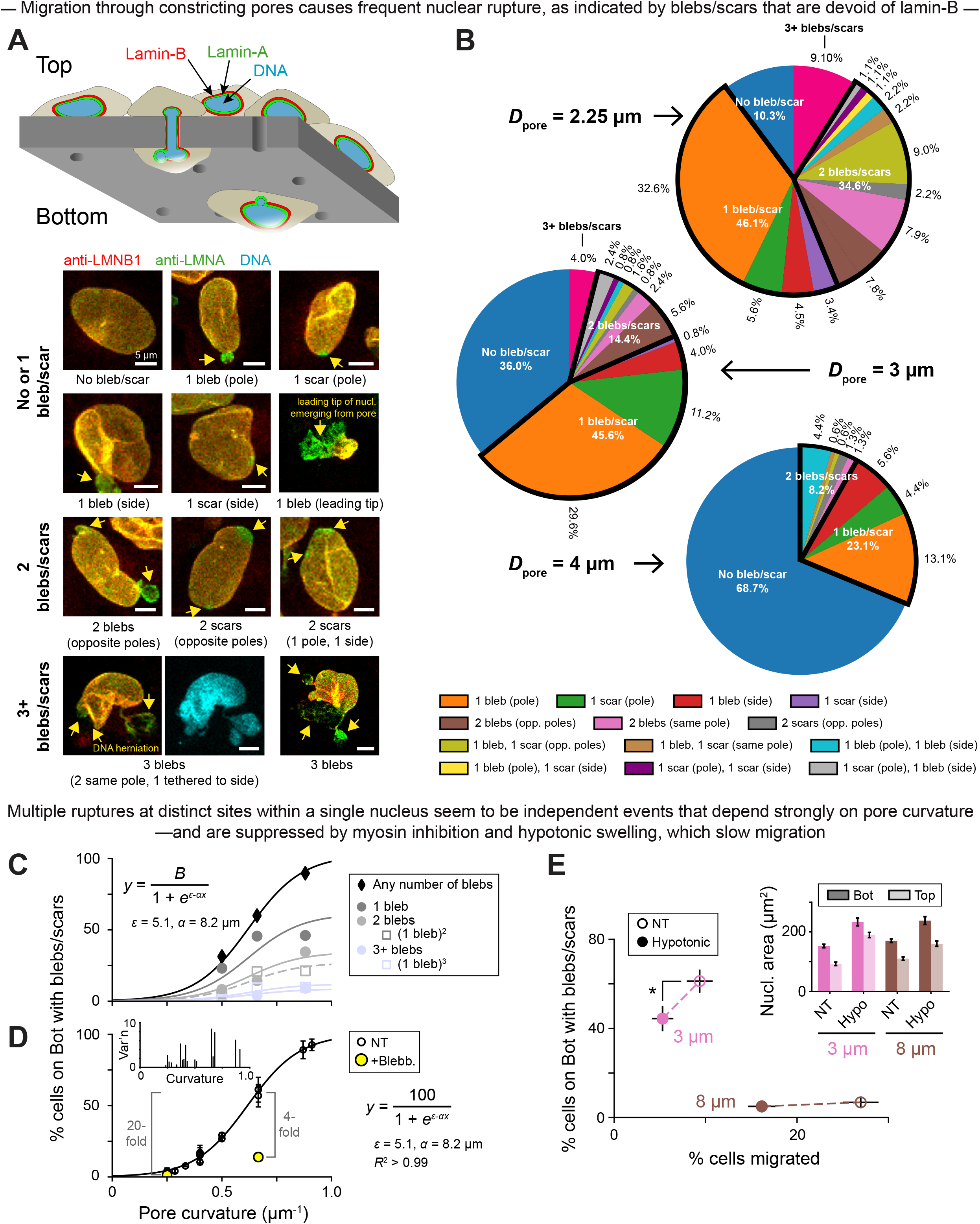
Migration through constricting pores causes frequent nuclear rupture; multiple such ruptures at distinct sites within a single nucleus seem to be independent events that depend strongly on pore curvature. A. Nuclei rupture as cells migrate through constricting pores from the Top to the Bottom of a Transwell. Images: nuclear blebs and scars (yellow arrows) in migrated U2OS human bone cancer cells on the Bottom of a 3 *μ*m pore membrane. Some nuclei show multi-site rupture, with 2 or 3+ blebs/scars. B. Distributions of bleb/scar number and location among U2OS cells that have migrated through pores of varying diameter *D*_pore_. ≥90 cells per *D*_pore_. C. Rupture frequency during migration, as indicated by % migrated U2OS cells with blebs/scars, increases with pore curvature. This trend holds overall (black diamonds) as well as among migrated cells with 1 (dark gray), 2 (light gray), and 3+ (light blue) rupture sites. Filled points are measured values from panel B. Unfilled are estimates of multi-site rupture frequency, obtained by treating each rupture as an independent event with probability equal to the measured % of 1-bleb/scar cells. Sigmoidal fits are per our simple filament (un)binding model; from top to bottom curve, fit parameter *B* = 102, 61, 34, 27, 12, 9. D. Across an expanded range of pore diameters (*D*_pore_ = 3 *μ*m to 8 *μ*m), rupture frequency depends strongly on pore curvature. Addition of blebb. to the Bottom of the pore membrane suppresses rupture but not as much as a ~2.5-fold curvature reduction. Data adapted from Xia, Pfeifer 2019; ≥50 (U2OS) cells per *D*_pore_ mean ± SEM. Inset: SEM values, from the main plot, are highest at intermediate pore curvatures. E. Hypotonic medium was applied to the Top and Bottom of 3 and 8 *μ*m pore membranes during U2OS cell migration. Non-treated (NT) = isotonic medium. 303 cells. Inset: hypotonic medium increases nuclear area across all conditions. 539 cells; ≥39 cells per condition; mean ± SEM.

Actomyosin inhibition — which should suppress nuclear pressure — is again less effective than low curvature in rescuing rupture (**Fig.3D**). Likewise, even when migrating cells were subjected to hypoosmotic stress, which caused nuclear swelling consistent with higher intranuclear pressure, rupture frequency *decreased* (**Fig.3E**, **Fig.S3A**), as did average bleb size (**Fig.S3B**). Multi-site rupture again seemed to involve independent events (**Fig.S3C**). Importantly, both hypoosmotic stress (**Fig.3E**) and actomyosin inhibition (Xia et al., 2019) cause slower migration to the bottom of the Transwell device, motivating a final study on nuclear strain rate.

### Rate-driven, curvature-dependent dilution of lamin-A and rupture

Slow curvature changes seem to result in little to no dilution of the supposedly viscous lamin-A meshwork (Fig.1A-C, Fig.S1B,C; **Fig.4A**). To further investigate the effects of strain rate on lamina dilution and nuclear rupture, U2OS cells expressing lamin-B1-GFP were detached, actin-depolymerized and pulled into micropipettes of varying diameter at either a slow or fast rate (**Fig.4B,C**). Regardless of aspiration rate, nuclei always ruptured in the highest-curvature small-diameter pipettes and rarely ruptured in the lowest-curvature pipettes (Fig.4C-ii), consistent with results for constricting pores (Fig.3C,D). This experiment also revealed an association between rapid curvature imposition (i.e. fast nuclear extension into the pipette), lamin-B dilution at the leading tip to some extent, and rupture (Fig.4C-ii), which lends credence to the observation that slow migration under hypoosmotic stress could suppress rupture.

**Figure 4.**
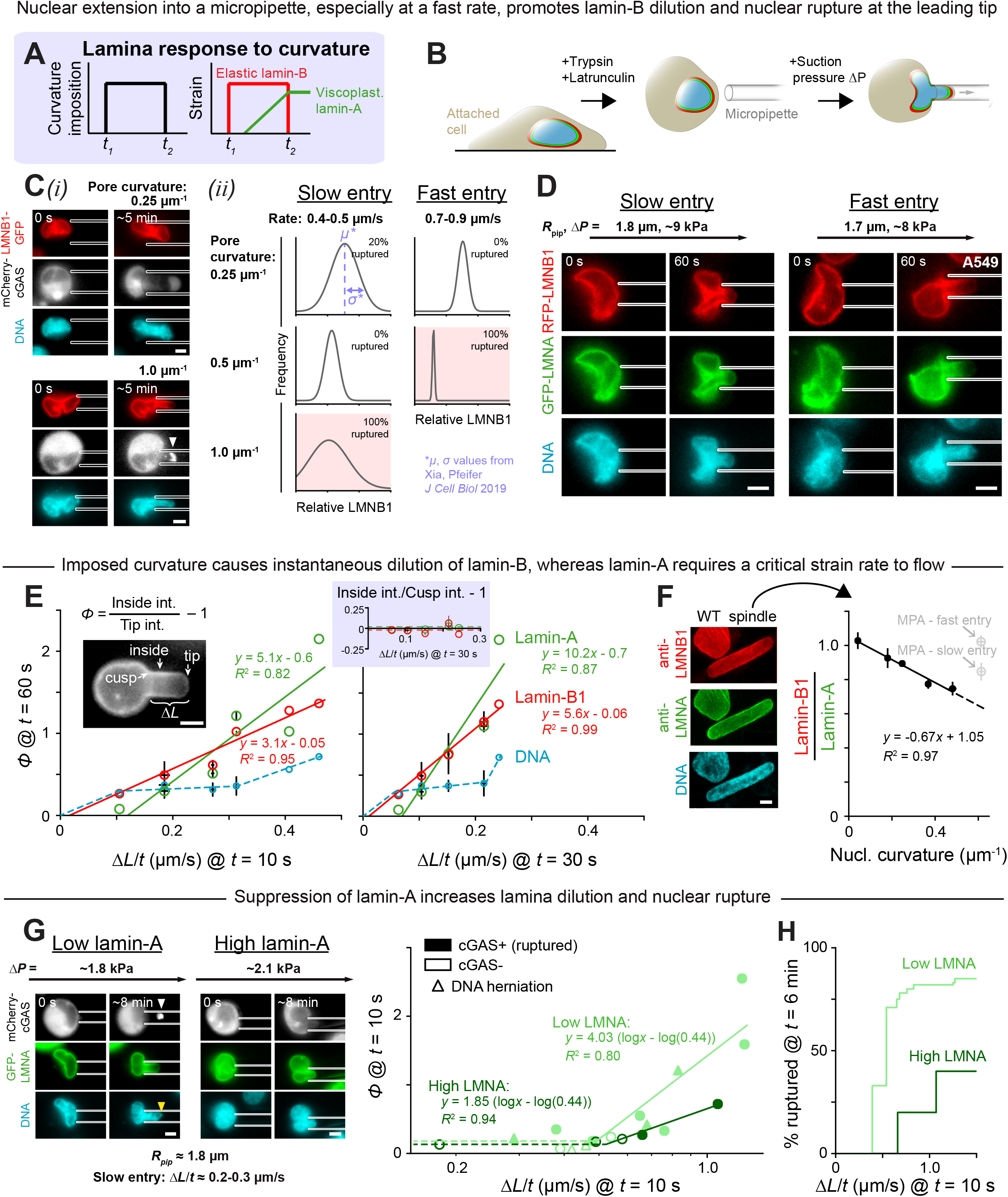
Nuclear entry into a constriction causes instantaneous dilution of lamin-B at the leading tip of the nucleus, while lamin-A requires a critical strain rate to flow. A. Lamin-A and lamin-B networks are predicted to have viscous and elastic responses, respectively, to applied stress. B. Cells were detached, treated with latrunculin to depolymerize the actin cytoskeleton, and pulled under controlled pressure Δ*P* into micropipettes. C. U2OS cells were transfected with lamin-B-GFP and mCherry-cGAS. (*i*) Representative images of cell aspiration into a low-curvature or a high-curvature pipette. Arrow head = nuclear envelope rupture, indicated by nuclear accumulation of cGAS. Scale = 5 *μ*m. (*ii*) For 17 cells pulled at a slow or fast rate into pipettes of varying curvature, lamin-B intensity at the nuclear tip was measured at *t* = 15 s. Plots show distributions of these intensities for all conditions. Based on mean μ ± SEM *σ* values from Xia, Pfeifer 2019; ≥3 cells per condition. D. A549 cells with gene-edited RFP-lamin-B were transfected with GFP-lamin-A and aspirated at varying rates into pipettes of fixed curvature ≈ 0.5 *μ*m^−1^ to examine resulting lamina dilution. *R*_pip_ = pipette radius. E. For 14 aspirated cells (panel D), nuclear tip dilution *ϕ* of lamin-A, lamin-B, and DNA was calculated (left inset) at *t* = 60 s. Plots show *ϕ* as a function of nuclear entry rate into the pipette Δ*L*/*t*, measured at an early timepoint (left) and slightly later timepoint (right). Right inset shows that, regardless of nuclear entry rate, lamins-A and -B do not dilute at the cusp at the pipette tip. 1-4 cells per bin; mean ± SEM for each bin. F. Images: select ‘spindle-shaped’ A549 clones have stably elongated nuclei as compared to wild-type (WT). Scale = 5 *μ*m. Plot: measurements taken at poles of spindle-shaped nuclei in 2D culture (black points) and at the leading tip of WT nuclei in micropipettes (gray points, data from panel E). 298 spindle-shaped cells; ≥11 cells per bin; mean ± SEM for each bin. G. U2OS cells with lamin-A knockdown were transfected with GFP-lamin-A—yielding a wide range of lamin-A levels—and mCherry-cGAS. Images: aspiration of a low-lamin-A and a high-lamin-A cell into a pipette of curvature ≈ 0.5 *μ*m^−1^. Arrow heads = nuclear envelope rupture, indicated by nuclear accumulation of cGAS (white) and DNA herniation (yellow). Scale = 5 *μ*m. Plot: nuclear tip dilution *ϕ* of lamin-A versus nuclear entry rate into the pipette Δ*L*/*t*. Solid lines are fits to the 11 low-lamin-A and 4 high-lamin-A cells with the highest Δ*L*/*t*; dashed lines guide the eye. cGAS+/- = does/does not exhibit nuclear cGAS accumulation. H. Low lamin-A cells rupture with enhanced frequency at a slow nuclear extension rate. Cells from panel G. % ruptured comprises cGAS+ cells and/or cells with DNA herniation.

The micropipette aspiration experiment was repeated with the endogenously tagged RFP-lamin-B1 A549 cells transfected with GFP-lamin-A. The cells were pulled at varying rates into pipettes of curvature ~0.5 *μ*m^−1^ (**Fig.4D,E, Fig.S4A**). As a nucleus extended into the pipette, lamin-B diluted instantaneously at the leading tip, proportional to extension rate Δ*L/t*, whereas lamin-A diluted only above a critical extension rate (Δ*L*/*t*)_crit_ at the same timepoint in aspiration (Fig.4E). At very long times, lamin-A diluted more like lamin-B and independent of initial rate (**Fig.S4B**). Notably, neither lamin-A nor -B diluted at the cusp created by the pipette wall (Fig. 4E), which suggests that both principal curvatures at a site must be the same sign (Fig. S1) — as at a nuclear pole — to cause lamin filament dissociation and dilution at that site. Because extension rates below (Δ*L*/*t*)_crit_ were difficult to probe with aspiration, we studied the ‘spindle-shaped’ A549 clones that exhibit a micropipette-like curvature but under slow, cell-generated forces in 2D culture (Fig.S1B,C): lamin-B dilutes at the highly curved nuclear poles while lamin-A does not (**Fig.4F**). The lamin-A layer thus requires a high strain rate to flow and dilute. This rate-dependent behavior is typical of viscous materials and might underlie the association of slow migration and reduced nuclear rupture in both hypotonically swollen and actomyosin-inhibited cells. Thus, when curvature imposition is slow, lamin-A dilutes minimally or not at all, allowing it to fortify the nuclear envelope against rupture even with lamin-B dilution. Consistent with such a mechanoprotective role for lamin-A, low-lamin-A cells exhibit greater lamina dilution and more frequent nuclear rupture when compared to high-lamin-A cells (**Fig.4G,H**, **Fig.S4C-E**).

Overall, our results support a model in which lamin-B filaments stably bind to low-curvature nuclear membranes but are too stiff to bend along high-curvature membranes—and thus detach quickly when curvature is applied. In comparison, Lamin-A requires a critical strain rate to flow: for slow curvature imposition, the lamin-A layer remains minimally deformed, but for rapid curvature imposition, lamin-A dilutes with lamin-B, and the envelope ruptures with high frequency. Interestingly, the method which shows that local, transient tensile stress on the nuclear membrane causes membrane rupture (Zhang et al., 2019), also causes high nuclear curvature at the rupture site. Likewise, across multiple primary cell types or established lines and with multiple experimental strategies here, actomyosin stress and intranuclear pressure appear to be weak modulators of lamina dilution and nuclear rupture compared to the strong responses to positive Gaussian curvature.

Lamin defective mice exhibit most or all of the basic tissue lineages (i.e. minimal differentiation defects) and are near-normal in size up to birth with some key tissue-specific proliferation defects. For example, lamin-B mutants show rupture and proliferation defects in brain (Vergnes et al., 2004), and because lamin-A is very low in soft brain, brain cells normally depend largely on lamin-B; stiffer tissues will have high lamin-A and be protected from lamin-B defects. Deeper physical insight into interphase rupture mechanisms thus motivates better understanding of lamin stoichiometry as well as lamin interactions (farnesylation, Lamin-B receptor, etc.). It is also foundational to clarifying effects on DNA damage, cell cycle disruption, and premature death of lamin defective organisms, from mice to man.

## Materials and methods

### Single-filament model

As previously described (Xia et al., 2019), we consider a single lamin-B filament that is stiff with a measurable length *L*_fil_ and persistence length ℓ_p_ (Turgay et al., 2017). The filament is either attached to or detached from the nuclear membrane; hence, in this ‘spherical cow’-level model, we write the following partition function for the filament:

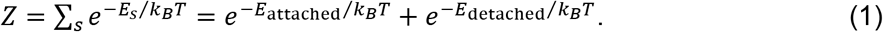

We take the detached state to be the reference state with *E*_detached_ = 0. If the nuclear membrane were flat, *E*_attached_ would simply be equal to a negative binding energy, −*E*, that favors filament attachment. But since the nuclear membrane can be curved (with curvature = 1/*R*), we consider two contributions to the energy of the attached state:

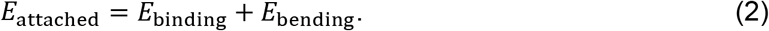

*E*_bending_ is the energy cost for bending the stiff filament along the curved membrane, and it is given by 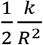, where *k* is the filament bending modulus. *E*_binding_ is not simply equal to −*E*, but is instead modulated by membrane curvature since curvature changes the filament-membrane contact area. Altogether, we can rewrite Eq. 2 as a function of curvature:

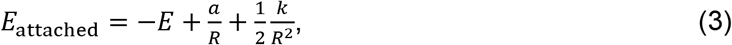

with *a/R*, where *a* is a constant, capturing the curvature-induced change in contact area between the nuclear membrane and the lamin-B filament. The second and third terms increase with 1/*R*, meaning that filament attachment becomes more energetically costly at high membrane curvatures—and detachment becomes more likely. The probability of the detached state is:

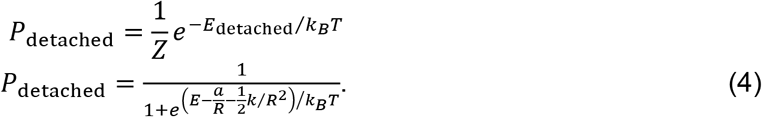

Indeed, in the high-curvature limit, 1/*R* becomes very large such that

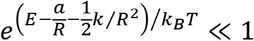

and *P*_detached_ → 1, consistent with high probability of lamin-B dilution and nuclear envelope rupture at the high-curvature poles of nuclei (Fig.1C, Fig.2D) and in small pores (Fig.3C,D) and pipettes (Fig.4C). Meanwhile, in the low-curvature limit, 1/*R* → 0, *P*_detached_ → 1/(1 + *e^E/k_B_T^*) → 0, assuming *E* ≫ *k_B_T*. That is, a lamin-B filament is unlikely to detach from a nuclear membrane of low curvature, which is consistent with low probability of lamin-B dilution and rupture on the flat sides of nuclei or in large pores and pipettes.

Note that the filament bending modulus *k* depends on filament length *L*_fil_ and persistence length ℓ_p_: *k* = (ℓ_p_*k_B_T*)*L*_fil_. We use the values *L*_fil_ = 0.38 *μ*m and ℓ_p_ = 0.5 *μ*m from (Turgay et al., 2017) to calculate *k* = (ℓ_p_*k_B_T*)*L*_fil_ = (0.5 *μ*m)(0.38 *μ*m)*k_B_T* = (0.19 *μ*m^2^)*k_B_T*. When fitting data (Fig.2D, Fig.3C,D), we find that the bending term 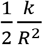 has almost no effect on the best-fit curve, so we exclude it. Fits are thus of the form

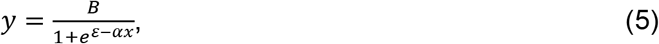

where *B* is a fit parameter and *ε* and *α* are, respectively, a binding energy and an interaction energy that were previously determined (Xia et al., 2019).

### Cancer cell lines

U2OS human osteosarcoma cells were a gift from Dr. Roger A. Greenberg of the University of Pennsylvania, Philadelphia, USA. A549 human lung adenocarcinoma cells with endogenously RFP-tagged lamin-B1 were purchased from MilliporeSigma (product number: CLL1149). U2OS and A549 cells were cultured in DMEM high glucose medium (Gibco) and Ham’s F12 medium (Gibco), respectively, and both were supplemented with 10% fetal bovine serum (FBS; MilliporeSigma) and 1% penicillin/streptomycin (Gibco). Cells were kept at 37°C and 5% CO_2_.

### Progeria MSCs

Primary fibroblast-derived induced pluripotent stem cells (iPSCs) were acquired from the Progeria Research Foundation Cell & Tissue Bank. iPSC differentiation into mesenchymal stem cells (MSCs) was performed as described in (Zou et al., 2013). Three days after initial splitting, iPS culture medium was replaced with MSC medium (low glucose DMEM supplemented with 10% FBS and 1% penicillin/streptomycin), which was then refreshed every two days. After two weeks of culture, cells were detached using 0.25% trypsin-EDTA (Gibco) and expanded on 0.1% gelatin coated dishes (BD Biosciences); cells were passaged whenever they reached confluence. After the third passage, a homogenous fibroblast-like cell population was achieved.

### Embryonic chick hearts

Embryonic chick hearts were extracted from White Leghorn eggs (Charles River Laboratories; Specific Pathogen Free (SPF) Fertilized eggs, Premium #10100326). SPF eggs were incubated at 37°C and 5% CO_2_ and rotated once per day for four days (to reach E4). Embryo removal was performed by windowing eggs and cutting major blood vessels to the embryonic disc tissue. Each extracted embryo was placed in a dish containing PBS, set on a 37°C-heated plate, and decapitated. The conotrunucus and sino venosus were severed to obtain the whole heart tube, which was then incubated at 37°C in pre-warmed chick heart media (α-MEM supplemented with 10% FBS and 1% penicillin/streptomycin) for at least 1 hour prior to use.

### Transfection/knockdown

GFP-lamin-A was a gift from Dr. David Gilbert of Florida State University, Tallahassee, USA (Izumi et al., 2000); GFP-KU80 was a gift from Dr. Stuart L. Rulten of the University of Sussex, Brighton, UK (Grundy et al., 2012); and mCherry-cGAS was a gift from Dr. Roger A. Greenberg of the University of Pennsylvania, Philadelphia, USA (Harding et al., 2017). We used small interfering RNAs (siRNAs) purchased from Dharmacon (ON-TARGETplusSMARTpool siLMNA, L-004978-00; and non-targeting siRNA, D-001810-10). For cells: we transfected either siLMNA (25 nM) or GFP/mCherry (0.5 ng/mL) with 1 *μ*g/mL Lipofectamine 2000 (Invitrogen, Life Technologies) for 72 hours (siLMNA) or 24 hours (GFP/mCherry) in corresponding media supplemented with 10% FBS. Knockdown efficiency of siLMNA was determined by immunofluorescence microscopy and by western blot via standard methods. For embryonic chick hearts: we transfected GFP and mcherry (0.2 – 0.5 ng/ml) as described above.

### Immunostaining

Cells were fixed in 4% paraformaldehyde (PFA; MilliporeSigma) for 15 minutes, permeabilized by 0.5% Triton-X 100 (MilliporeSigma) for 10 minutes, and blocked with 5% bovine serum albumin (BSA; MilliporeSigma) for 30 minutes. Cells were then incubated in primary antibodies overnight at 4°C. Primary antibodies used were lamin-A/C (1:500, mouse, Cell Signaling #4777S) and lamin-B1 (1:500, rabbit, Abcam ab16048). The next day, cells were incubated in secondary antibodies (1:500, donkey anti-mouse or -rabbit, Thermo Fisher) for 90 minutes before DNA was stained using 8 *μ*m Hoechst 33342 (Thermo Fisher) for 15 minutes. Unless otherwise specified, steps were carried out at room temperature with gentle agitation by orbital shaker. ProLong Gold Antifade (Invitrogen, Life Technologies) was used as a mountant. Embryonic chick hearts were fixed in 4% PFA for 45 minutes, washed using PBS, and permeabilized using 0.5% Triton-X (Fisher) for 30 minutes. The hearts were incubated in 16 *μ*m Hoescht for 2 hours prior to being mounted between two glass coverslips.

### Imaging

Confocal imaging of fixed cells and tissues was done on a Leica TCS SP8 system with 63×/1.4-NA oil-immersion and 40×/1.2-NA water-immersion objectives. Live imaging of cells and tissues was performed under normal culture conditions (37°C and 5% CO_2_, complete medium). Cells were live-imaged using either an EVOS FL Auto Imager (Fig.1B,C) or a Zeiss Axio Observer 7 (Fig.1D), both with 40× objectives; embryonic chick hearts were live-imaged using an Olympus I81 with a 4× objective and a CCD camera. Epifluorescence imaging of fixed progeria cells was obtained using an Olympus IX71 with a digital EMCCD camera equipped with a 40×/0.6-NA objective.

### Image analysis

Image analysis was performed in ImageJ (Schneider et al., 2012).

#### Measuring lamin-B-to-A ratio

To measure lamin-B-to-A ratio at a nuclear pole, lamin-B and lamin-A intensity profiles were generated along the nucleus’s major axis, originating at the pole of interest (e.g. Fig.1B). Each intensity profile was min-max normalized, as follows:

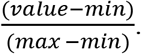

From these plots, we computed average lamin-B and average lamin-A intensity within a ~1 *μ*m distance of the pole (i.e. from *x* = 0 to *x* ≈ 1 *μ*m in the plots), and then took the ratio of those two values. We used the same approach to measure lamin-B-to-A ratio at nuclear sides, except intensity profiles were generated along the minor rather than major axis.

#### Measuring curvature

At the point of interest along the inner nuclear surface, an osculating circle—or “circle of best fit”—was drawn to overlap the nuclear surface for a distance of 2 *μ*m into the cell. Curvature was then calculated as 1/*R*, where *R* is the radius of the osculating circle. Similarly, the curvature of a given pore/pipette was taken to be the inverse of the pore/pipette radius.

#### Identifying nuclear blebs

To identify nuclear blebs/scars in a given image, we superimposed lamin-A fluorescence signal (green) on top of lamin-B fluorescence signal (red) such that nuclear regions of high-lamin-A/low-lamin-B (i.e. high-green/low-red) were easily detected by eye. For the progeria stem cells, which exhibit very weak staining by lamin-A/C antibodies, local loss of lamin-B was the sole criterion used to identify blebs/scars.

#### Measuring lamina dilution in micropipettes

To quantify lamin(-A or -B) dilution at the leading tip of an aspirated nucleus, we measured the average background-subtracted lamin intensities of the ‘tip’ and ‘inside’ regions (Fig.4E, inset). Intensities were measured within boxes of area approximating the pipette diameter squared. Lamin dilution was then calculated as the ratio of ‘inside’ intensity to ‘tip’ intensity—minus 1 so that dilution = 0 if the leading tip shows zero lamin loss. ‘Relative LMNB1’ (Fig.4C-ii) was calculated as simply the ratio of ‘tip’ intensity to ‘inside’ intensity.

### Pore migration

Cells were seeded at a density of 4.5 × 10^5^ cells/cm^2^ on commercially available polycarbonate filter membranes (Corning) with pore diameters of 3, 5, or 8 *μ*m and on etched membranes with modified pore diameters of 2.25, 4, 6, or 7 *μ*m. Pore etching was performed as previously described (Deviri et al., 2019; Xia et al., 2019). Complete medium was added to the Top and Bottom of each membrane such that no nutrient gradient was established; for the hypoosmotic stress experiments, complete medium was replaced by a 2:3 mixture of high glucose DMEM and ddH_2_O, supplemented with 10% FBS and 1% penicillin/streptomycin. Cells were allowed to migrate from membrane Top to Bottom over the course of 24 hours under normal culture conditions (37°C and 5% CO_2_). After the 24-hour migration period, each membrane—with un-migrated cells attached on Top and migrated cells attached on Bottom—was fixed, stained, and imaged by confocal microscopy as described above (Immunostaining, Imaging). Membranes were excised from their plastic inserts prior to cell permeabilization.

### Micropipette aspiration

To prepare for aspiration, cells were detached using 0.05% Trypsin-EDTA (Gibco), and incubated in 0.5 μg/mL latrunculin-A (MilliporeSigma) and 8 *μ*m Hoechst 33342 for 30 minutes at 37°C. During aspiration, cells were resuspended in PBS with 1% BSA and 0.2 μg/mL latrunculin-A. Pipette diameters varied from ~2-8 *μ*m, and aspiration pressures varied from ~1-10 kPa; see figures and figure legends for the parameters of each experiment. Aspiration was monitored over ~5-60 minutes and imaged using a Nikon TE300 with a 60×/1.25-NA oil-immersion objective and a digital EMCCD camera (Photometrics).

### Drug perturbations

To image cells undergoing mitosis, we synchronized A549 cells at G2- or M-phase by treating with 50 nM nocodazole for 16 hours. Nocodazole was then gently washed out with pre-warmed PBS ×5, allowing cells to proceed through mitosis. To inhibit myosin II, progeria stem cells and embryonic chick hearts were treated with 25 *μ*m blebbistatin for 2 hours prior to fixation and staining. In the pore migration assay with myosin II inhibition, 20 *μ*m blebbistatin was added to the Bottom of the filter membrane for the entire migration period. In every case, controls were treated with an equal concentration of vehicle solvent DMSO.

### Statistics

All statistical analyses were conducted using Microsoft Excel and Python. Statistical comparisons were made by unpaired two-tailed Student’s *t*-test and were considered significant if *p* < 0.05. Unless mentioned specified, *n* indicates the number of samples, cells, or wells quantified in each experiment.

## Acknowledgements

We thank the Penn Cell & Developmental Biology Microscopy Core’s Dr. Andrea Stout and Jasmine Zhao for microscope access and expert technical assistance. We thank Larry Dooling for developing the pore etching method that enabled the custom-made transwell pores.

## Author contributions

C.R.P and M.P.T. designed and performed all experiments except those noted below, analyzed data, made figures, and co-wrote the manuscript. S.C. and M.V. designed and performed the embryonic chick heart experiments. L.L.V., E.G.R.-D., and K.T.S. performed the hypotonic swelling experiments, and L.J.D. made the custom-sized pores. D.E.D. helped design experiments, analyzed data, made figures, conceived of the single filament model, and co-wrote the manuscript.

## Supplemental figure legends

**Figure S1.**
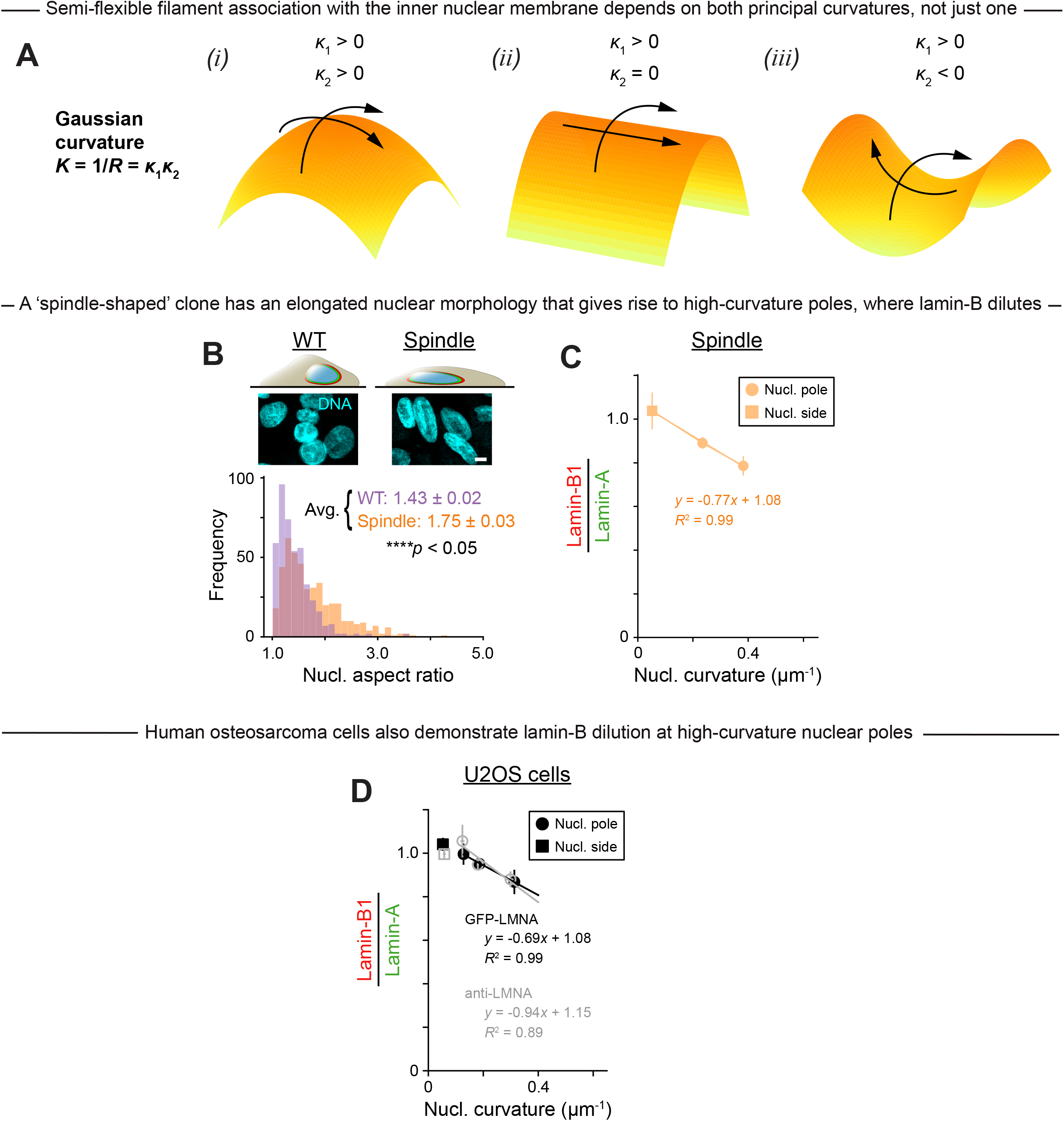
Lamin-B filaments tend to dissociate from nuclear regions of high Gaussian curvature, such as the poles of stably elongated nuclei in ‘spindle-shaped’ cells. A. The Gaussian curvature *K* of a surface at a given point is the product of the principal curvatures, *κ*_1_ and *κ*_2_, at that point. Three surfaces illustrate (i) positive, (ii) zero, and (iii) negative Gaussian curvature. For a nuclear surface to induce lamin-B detachment, both principal curvatures must be high/non-zero, as in (i) and (iii). Gaussian curvature can also be expressed in terms of the Gaussian radius of curvature *R*: *K* = 1/*R*. B. In normal 2D culture, some A549 RFP-lamin-B cells have an elongated morphology similar to that of mesenchymal stems cells. These ‘spindle-shaped’ A549 cells, which were cloned for further study, have elongated nuclei—with an average aspect ratio significantly higher than bulk wild-type (WT) A549 nuclei. ≥439 cells per condition; mean ± SEM; scale = 5 *μ*m. C. Lamin-A and -B intensities were measured at the poles and sides of ‘spindle-shaped’ A549 nuclei. Plot shows that lamin-B depletes relative to lamin-A as nuclear curvature increases, suggesting that high curvature, slowly imposed, is sufficient to dilute lamin-B but not lamin-A. 91 cells; ≥10 cells per bin. D. Lamin-B dilution at high-curvature nuclear poles was confirmed using antibodies against lamin-A and lamin-B (black) or transfected GFP-lamin-A and antibody-stained lamin-B (gray) in U2OS cells. The trend is the same whether the assay was performed with immunostained or fluorescently tagged proteins. 13 cells per condition for nuclear sides; ≥59 cells per condition (≥5 cells per bin) for nuclear poles; mean ± SEM.

**Figure S2.**
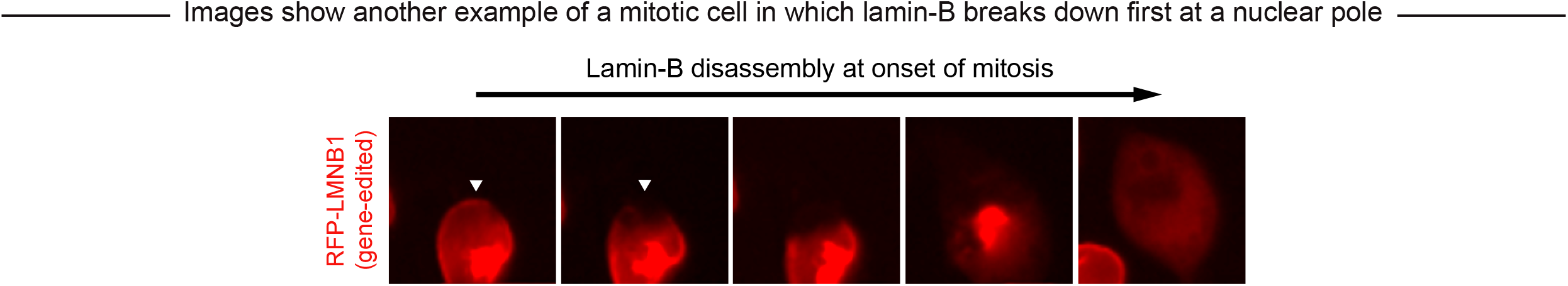
Nuclear curvature might contribute, along with other mechanisms, to lamin-B disassembly during mitosis. See Fig.1D. Live images of an A549 cell undergoing mitosis show the lamin-B network breaking down first at the pole of the nucleus (arrow heads).

**Figure S3.**
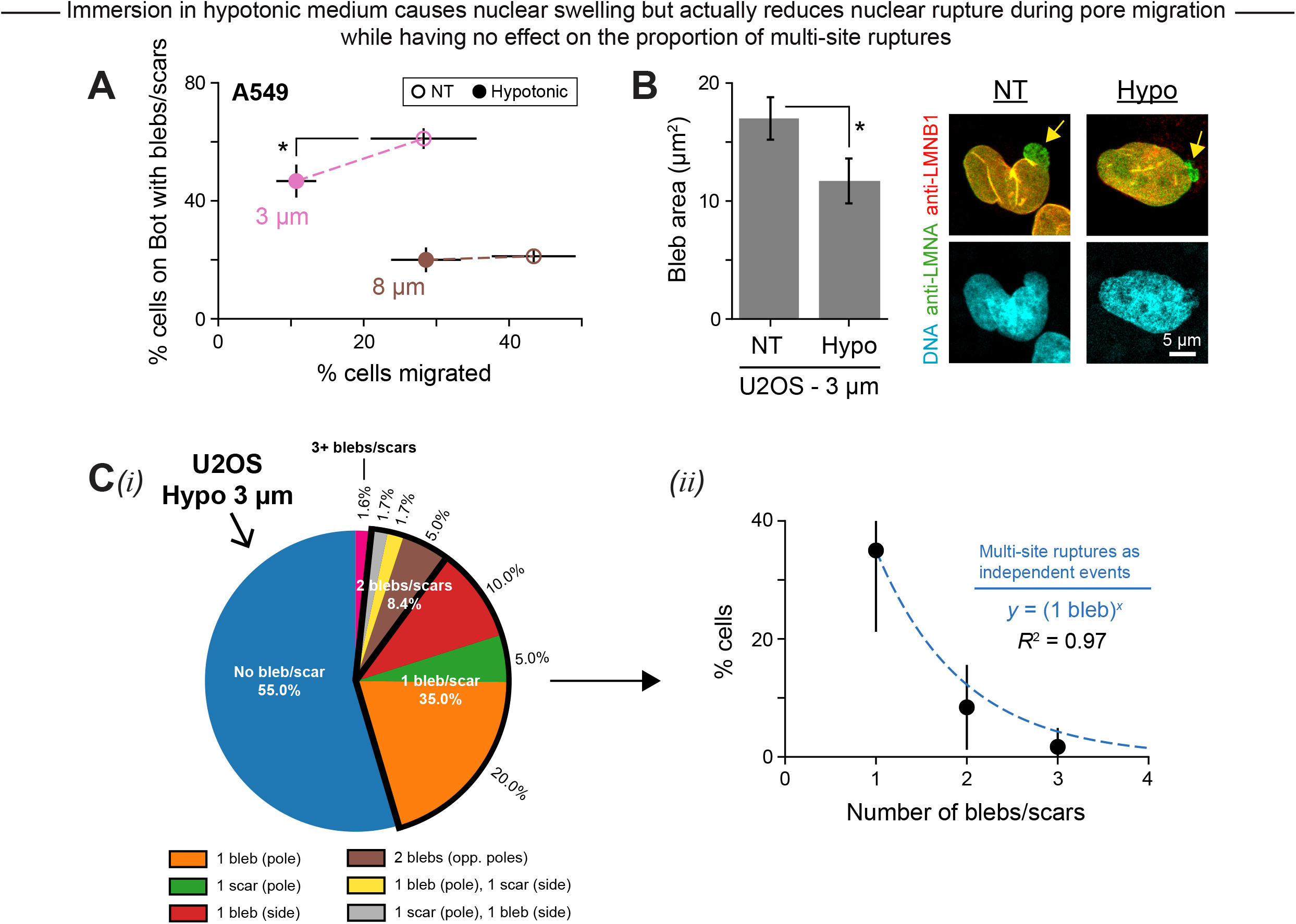
Hypoosmotic stress slows migration through constricting pores and reduces migration-induced rupture overall—but multi-site rupture still occurs with predictable frequency. A. Same as Fig.3E, except with A549 cells. Hypotonic medium was applied to the Top and Bottom of 3 and 8 *μ*m pore membranes during A549 cell migration. 448 cells; ≥91 cells per condition; mean ± SEM. B. See Fig.3E. Plot: In addition to decreasing nuclear rupture frequency, hypotonic medium also decreased nuclear bleb size among 3 *μ*m pore-migrated U2OS cells. Images: U2OS cell nuclei after migration. ≥28 blebs per condition; mean ± SEM. C. See Fig.3A-C. (*i*) Distribution of bleb/scar number and location among U2OS cells that migrated through 3 *μ*m pores under hypotonic conditions. 60 cells. (*ii*) Plot shows the % of these cells with 1, 2, and 3 blebs/scars. If each nuclear rupture—and thus each bleb/scar formation—is an independent event, then the probability of *x* blebs/scars co-occurring in a single nucleus is (1 bleb)^*x*^, where (1 bleb) is the probability of 1 bleb/scar occuring (i.e. the measured % of 1-bleb/scar cells). The function *y* = (1 bleb)^*x*^ fits well to data, suggesting that multiple ruptures at distinct sites within a single nucleus are indeed independent events.

**Figure S4.**
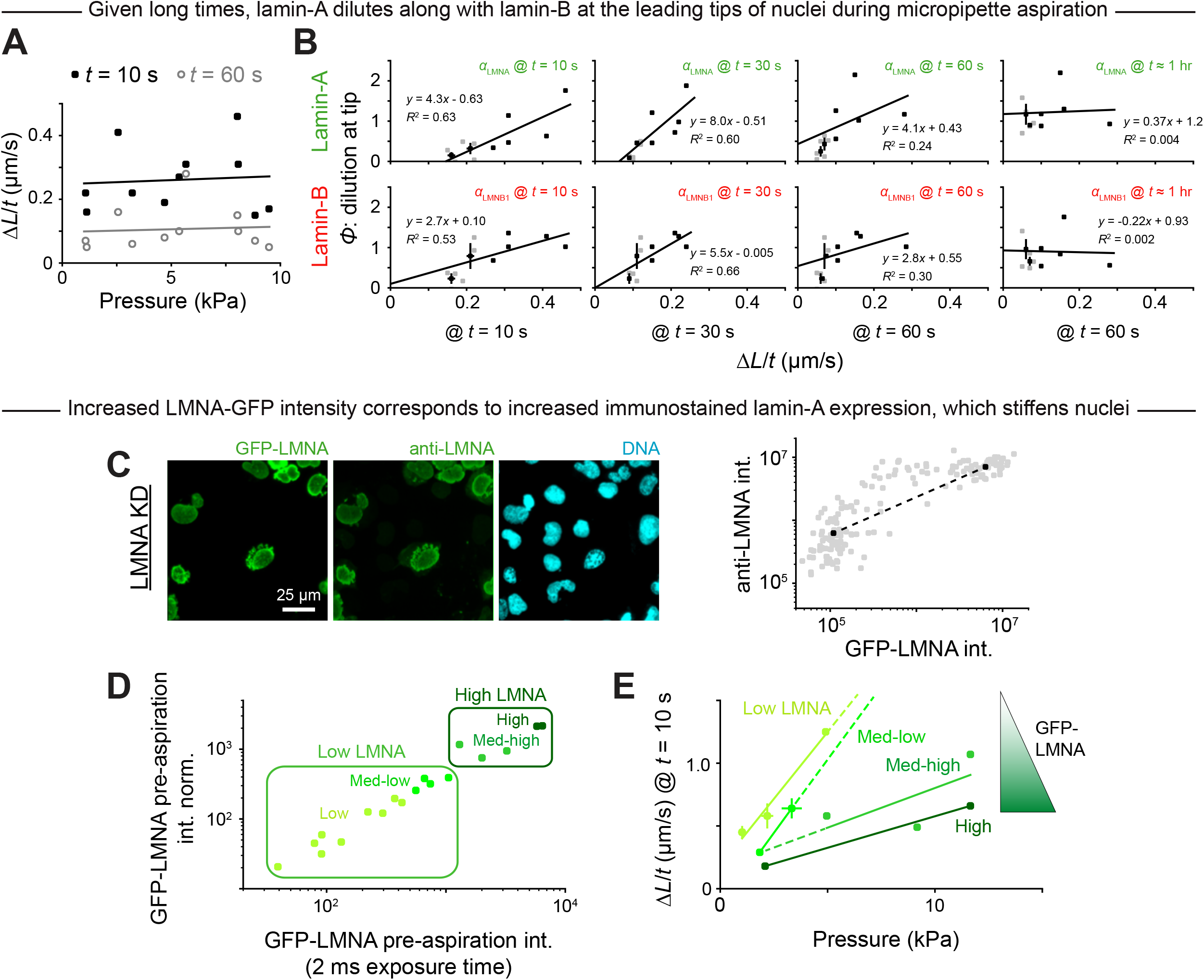
Lamin-A exhibits rate- and time-dependent flow, and rescue of lamin-A knockdown with over-expression of GFP-lamin-A yields a wide range of lamin-A levels and corresponding nuclear stiffnesses. A. See Fig.4D,E. A549 RFP-lamin-B cells were transfected with GFP-lamin-A and aspirated at varying rates into pipettes of fixed curvature ≈ 0.5 *μ*m^−1^. Nuclear extension rate Δ*L/t* correlated poorly with applied pressure at early (black) and slightly later (gray) timepoints, improving confidence that lamina dilution under rapid nuclear extension is truly a rate effect rather than an applied pressure effect. 14 cells. B. Cells from panel A. Nuclear tip dilution *ϕ* of lamin-A and lamin-B is plotted against rate of nuclear extension into the pipette Δ*L*/*t*, measured at various timepoints in aspiration. A critical strain rate, required for lamin-A to flow, is evident at the early timepoints. Given long times (t ≈ 1 hour), most aspirated cells showed both lamin-A and lamin-B dilution at the leading tip of the nucleus regardless of initial extension rate. C. Images: U2OS cells with lamin-A knockdown (KD) were transfected with GFP-lamin-A, and then fixed and stained with antibodies against lamin-A. Plot: in these KD cells, increasing GFP-lamin-A intensity corresponds to increasing lamin-A expression based on anti-lamin-A staining. Dashed line is meant to guide the eye. 168 cells. D. See Fig.4G,H. We performed micropipette aspiration of 18 lamin-A knockdown U2OS cells that were transfected with GFP-lamin-A and mCherry-cGAS. Prior to aspiration, the cells exhibited a wide range of GFP-lamin-A intensities, as confirmed by two different but highly correlated metrics: intensity norm. to optimized exposure time (=20-600 ms) and intensity at a set exposure time of 2 ms. E. Cells from panel D. When pulled into pipettes of curvature ≈ 0.5 *μ*m^−1^, low-lamin-A nuclei were found to be more compliant than high-lamin-A nuclei, with higher rates of nuclear extension Δ*L/t* at a given aspiration pressure. 1-4 cells per bin.

